# 3D spheroid culturing of *Astyanax mexicanus* liver-derived cell lines recapitulates distinct transcriptomic and metabolic states of *in vivo* tissue environment

**DOI:** 10.1101/2023.06.09.544423

**Authors:** Tathagata Biswas, Naresh Rajendran, Huzaifa Hassan, Chongbei Zhao, Nicolas Rohner

## Abstract

*In vitro* assays are crucial tools for gaining detailed insights into various biological processes, including metabolism. Cave morphs of the river-dwelling fish species, *Astyanax mexicanus*, have adapted their metabolism allowing them to thrive in the biodiversity-deprived and nutrient-limited environment of caves. Liver-derived cells from the cave and river morphs of *Astyanax mexicanus* have proven to be excellent *in vitro* resources to better understand the unique metabolism of these fish. However, the current 2D cultures have not fully captured the complex metabolic profile of the *Astyanax* liver. It is known that 3D culturing can modulate the transcriptomic state of cells when compared to its 2D monolayer culture. Therefore, in order to broaden the possibilities of the *in vitro* system by modeling a wider gamut of metabolic pathways, we cultured the liver-derived *Astyanax* cells of both surface and cavefish into 3D spheroids. We successfully established 3D cultures at various cell seeding densities for several weeks and characterized the resultant transcriptomic and metabolic variations. We found that the 3D cultured *Astyanax* cells represent a wider range of metabolic pathways, including cell cycle changes and antioxidant activities, associated with liver functioning as compared to its monolayer culture. Additionally, the spheroids also exhibited surface and cave-specific metabolic signatures, making it a suitable system for evolutionary studies associated with cave adaptation. Taken together, the liver-derived spheroids prove to be a promising *in vitro* model for widening our understanding of metabolism in *Astyanax mexicanus* and of vertebrates in general.

## 1. Introduction

Different populations of the fish species *Astyanax mexicanus* have proven to be valuable models for studying metabolic adaptation under nutrient-limited conditions. The species exists in two major morphs –river dwelling surface fish, and cave-adapted cavefish. Despite the nutrient-deficient cave environment, cavefish have evolved various metabolic adaptations to survive and thrive. These adaptations include hyperphagia, high blood sugar, enhanced adipogenesis, improved glycogen production, and efficient recycling of amino acids (Aspiras, Rohner, Martineau, Borowsky, & Tabin, 2015; Krishnan, Seidel, et al., 2022; Olsen et al., 2023; Riddle et al., 2018; Xiong, Krishnan, Peuss, & Rohner, 2018). Therefore, comparative studies between the metabolic adaptations of cavefish and the typical metabolism of their river-dwelling cousins provide a tractable opportunity to gain organism-level insight of metabolism in a living animal model.

Nevertheless, the relatively lengthy generation time of at least 5 months (Kozol et al., 2023) and other husbandry constraints have made high-throughput approaches challenging. In order to overcome these challenges and to provide a more focused attempt at dissecting metabolic pathways associated with specific cell types, we realized the importance of in vitro and explant based systems (Pagano et al., 2018; Torres-Paz & Retaux, 2021), and recently generated liver-derived stable cell lines from both the cave and surface morphs (Krishnan, Wang, et al., 2022; Rajendran et al., 2023). These cell lines, termed as <u>C</u>ave<u>F</u>ish <u>L</u>iver-derived (CFL) and <u>S</u>urface <u>F</u>ish <u>L</u>iver-derived (SFL), express liver-specific enzymes and efficiently capture key metabolic traits of cavefish adaptation (Krishnan, Wang, et al., 2022). Yet, the cell line transcriptomes indicate that the liver-derived cell lines do not fully replicate the *in vivo* transcriptomes of cavefish or surface fish livers.

It is known that 3D culturing of cells *in vitro* significantly changes the metabolic as well as the associated transcriptomic state of the system, compared to their 2D cultured counterparts (Lagies et al., 2020; Park et al., 2022; Rybkowska et al., 2023; Tekin et al., 2018). Therefore, we reasoned that culturing liver-derived cells into 3D spheroids could activate more metabolically relevant pathways presenting an opportunity to study these pathways *in vitro*. In this study, we examined the response of liver-derived *Astyanax* cells upon 3D culturing into viable spheroids. Both SFL and CFL 3D spheroids were cultured for up to four weeks at varying seeding densities. Upon successful culturing of both SFL and CFL spheroids, we characterized their transcriptomes and the resulting metabolic profiles. We found that 3D culturing elicits a common set of transcriptomic responses, in particular a decrease in cell cycle markers and an increase in antioxidant capacities. Along with the common set of responses, we also observed morph-specific responses in CFL and SFL, including enrichment for biological processes associated with response to xenobiotic stimulus and immune response, respectively. Taken together, we successfully cultured 3D spheroids from *Astyanax* cells, which would be an essential step towards understanding metabolic adaptation in cavefish and modeling more facets of metabolic diseases and challenges associated with liver functioning.

## 2. Materials and Methods

### 2.1. Generation of *Astyanax* liver-derived cell spheroids

For the generation of 3D spheroids, cells maintained in 2D (Krishnan, Wang, et al., 2022) were washed once with phosphate-buffered saline and then incubated with 0.25% trypsin–EDTA for 5 min at room temperature. The disassociated cells were suspended with complete media and centrifuged at 1000 rpm for 5 min. The composition of the complete media for *Astyanax* liver-derived cells was defined previously (Rajendran et al., 2023). The pelleted cells were re-suspended in complete media and counted using 0.4% trypan blue (1:1) and an automatic hemocytometer. Cells were seeded at different densities (10k; 25k; 50k & 100k cells/well) in ultra-low-attachment 96-well round-bottomed plates for 3D spheroid culture with 100μL complete media per well. Cells were maintained at 27°C without CO_2_. On day 3, the plates were placed in an orbital shaker at 70 rpm and half the amount of media was replaced every 4 days until the end of the experiment. The morphologies of the spheroids were captured at various time points using an inverted light microscope equipped with a digital camera (Leica DM-IL MC170).

### 2.2. RNA extraction, library preparation, and RNA sequencing

To collect total RNA from the spheroid samples, all 8 variants of spheroids (4 different densities of SFL and CFL each) were cultivated in 16 separate U-bottomed wells as biological replicates. Each set of densities from each morph was collected separately in 1.5ml centrifuge tubes and spun at 1000g for 2mins. After discarding the media, the samples were flash-frozen. Once all samples were gathered and frozen, total RNA was extracted using the *mir*Vana RNA isolation kit (Catalog number: AM1560) according to the manufacturer’s instructions. The mRNA-seq libraries were generated from 100ng of high-quality total RNA, as assessed using the Bioanalyzer (Agilent). Libraries were made according to the manufacturer’s directions for the TruSeq Stranded mRNA Library Prep kit (48 Samples) (Illumina, Cat. No. 20020594), and TruSeq RNA Single Indexes Sets A and B (Illumina Cat. No. 20020492 and 20020493). Resulting short fragment libraries were checked for quality and quantity using the Bioanalyzer (Agilent) and Qubit Fluorometer (Life Technologies). Libraries were pooled, requantified, and sequenced as 75bp single reads on a high-output flow cell using the Illumina NextSeq 500 instrument. Following sequencing, Illumina Primary Analysis version RTA 2.11.3.0 and bcl-convert-3.10.5 were run to demultiplex reads for all libraries and generate FASTQ files.

### 2.3. Correlation scatter plots, Heatmaps and GO analysis

To assess the correlation between different samples, scatter plots were generated using the R package ‘ggpubr’ (v0.4.0), based on gene expression values in ‘TPM’ (Transcripts per Million). p-values and *R*^2^ were computed using Pearson’s correlation.

Heatmaps were generated using R package ‘pheatmap’ (version 1.0.12). Gene expression values in TPM were used and genes having zero expression across all samples were excluded. The rows were scaled by computing the z-score for each gene across all samples.

### 2.4. Differential expression and gene ontology (GO) analysis

RNA-seq reads were demultiplexed into FASTQ format allowing up to one mismatch using Illumina bcl-convert (v3.10.5). Subsequently, the reads were aligned to *Astyanax mexicanus* reference genome from University of California at Santa Cruz (UCSC) using STAR (v2.7.3a) (Dobin et al., 2013). The gene model retrieved from Ensembl, release 102 was used to generate gene read counts. The transcript abundance ‘TPM’ (Transcript per Million) was quantified using RSEM (v1.3) (Li & Dewey, 2011). Differentially expressed genes were determined using R package edgeR (v3.38.4) (Robinson, McCarthy, & Smyth, 2010). Prior to differential expression analysis, low-expression genes were filtered out based on a cutoff of 0.5 CPM (Counts Per Million) in at least one library. The resulting p-values were adjusted with Benjamini-Hochberg method using R function p.adjust. Genes with an adjusted p-value < 0.05 and a fold change of 2 were considered as differentially expressed.

Gene functional enrichment analysis or gene ontology (GO) analysis was performed using a custom script built over R package ‘clusterProfiler’ (v4.4.4) (Wu et al., 2021). *Astyanax mexicanus* Gene-GO terms retrieved from Ensembl BioMart were used to identify over-represented GO terms in the differentially expressed genes compared to the background list of all genes.

### 2.4 Histology

Fish liver spheroids were fixed in 4% solution of paraformaldehyde (PFA) in PBS for 1hr at RT, followed by rinsing in PBS. Then samples were dehydrated through sucrose dehydration: first the PBS was replaced with 15% sucrose and incubated until the fish liver spheroids sunk to the bottom, then the 15% sucrose was replaced with 30% sucrose and incubated until fish liver spheroids sunk to the bottom. Dehydrated fish liver spheroids were embedded in OCT and subjected to cryo sectioning (10 μm). Sections were then stained in 5ug/ml WGA (Invitrogen™, Cat.: W11261) solution in PBS containing 10ug/ml DAPI (Bittium, Cat.: 40011) for 30 min. After rinsing out unbound WGA and DAPI with PBS, samples were mounted by Prolong Gold antifade reagent (life technologies, Cat.: P36930), and subjected to imaging under a fluorescence microscope (Zeiss Axiovert inverted fluorescence microscope).

### 2.8 Antioxidant Assay

The antioxidant assay was performed using the Antioxidant Assay Kit from Cayman Chemical (Item No. 709001) according to the manufacturer’s instructions and absorbance was measured at 405nm. For this assay, spheroids of each morph and seeding density were grown in replicates of 24 samples. For comparison to 2D culture, cells were also cultured in a 96-well flat-bottomed plate for 2D monolayer cell culture in similar numbers, i.e., 24 samples from each morph and density. The monolayer cells were cultured (Krishnan, Wang, et al., 2022) for 24hrs to minimize the chances of cell number variation due to proliferation. The spheroids were cultured for 14 days.

## 3. Results

### 3.1. Generation of 3D spheroids from liver-derived cell lines

To generate 3D spheroids from the liver-derived cells of *Astyanax*, we cultured the cells in U-bottomed ultra-low-attachment plates and let the cells self-aggregate (Park et al., 2022). To evaluate the dynamics of spheroid formation, we initially seeded 10,000 (10k) cells from both the surface fish liver-derived cell lines (SFL) and the cavefish liver-derived cell lines (CFL) into 3D cultures. Within a brief span of 24hrs, both the SFL and CFL cells were able to self-aggregate and form 3D spheroids (Figure 1a). To better assess long-term aggregation, we cultured SFL and CFL spheroids for a total of 4 weeks prior to characterization. Upon 4 weeks of culture, conspicuous spheroids persisted, similar to the ones formed at 24hrs (Supporting information Figure S1). Histological sections confirmed an overall healthy morphology and membrane markers exhibited uniform distribution of cells in the spheroids derived from both morphs (Figure 1b, c). It is of interest to note that SFL spheroids showed visibly lower affinity to dense aggregation as compared to the CFL-derived spheroids, as is evident from both the overall morphologies and histological sections (Figure 1b, c, S1). This observation aligns with previous observations showing that CFL cells express higher levels of cell adhesion molecules compared to SFL cells, potentially influencing cell adhesion and aggregation in 3D culture (Krishnan, Wang, et al., 2022).

**Figure 1:**
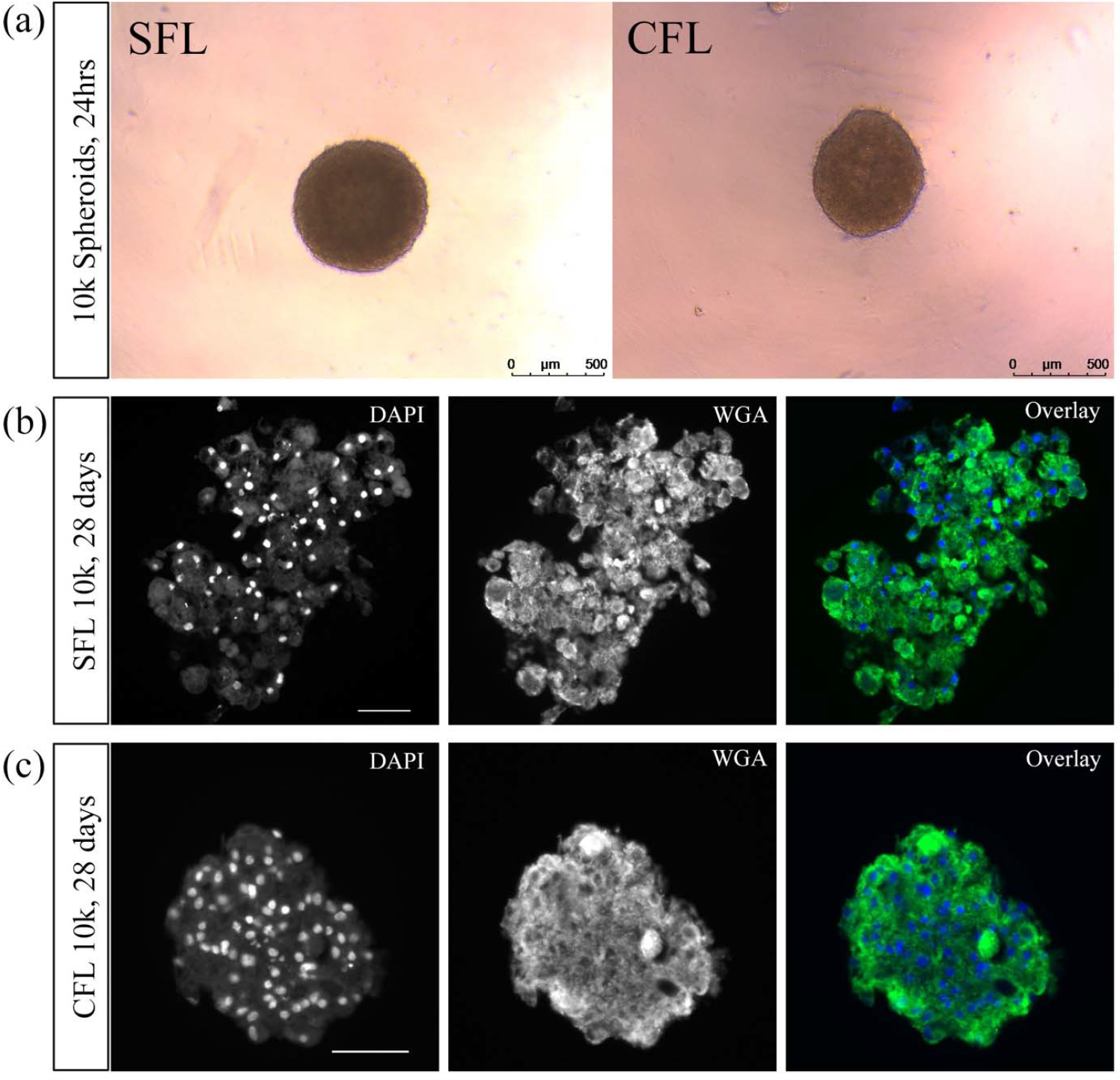
Spheroids from liver-derived cells of *Astyanax mexicanus*. (a) Spheroids from SFL and CFL developed within 24hrs of being 3D cultured in ultra-low attachment U-bottomed plates. Histological sections of both SFL (b) and CFL (c) spheroids after 4 weeks of culture. DAPI (blue) marks the nucleus and WGA (green) stains the cell membrane, scale-bar:50μm.

### 3.2. Altered transcriptomic profile of *Astyanax* cells upon spheroid formation

After successfully generating spheroids from SFL and CFL cells, we sought to understand how culturing *Astyanax* cells into 3D spheroids alters their transcriptomic profiles and whether varying cell seeding densities would elicit different responses. To this end, we used a series of increasing seeding densities of 10k, 25k, 50k, and 100k cells for generating spheroids from SFL and CFL cells. With 4 different seeding densities, we grew 8 different spheroid variants, 4 each for SFL and CFL cells. After 4 weeks of culturing, we extracted total RNA from these samples and performed bulk-RNAseq. In order to measure how culturing 3D spheroids changes transcription compared to the 2D cell cultures, we created scatter plots for both conditions and both types of cell lines. We used the existing 2D transcriptomic profile from our previous study for comparisons (Krishnan, Wang, et al., 2022). The correlation coefficients (*R*^2^) for SFL, ranged from 0.69 to 0.74, and that for CFL ranged from 0.68 to 0.72 across the varying seeding densities (Figure 2a-d and Supporting information Figure S2). These *R*^2^ values indicate that the overall transcriptomic profile indeed changes when transitioning from 2D to 3D cultures for both SFL and CFL cells. Further, gene expression heatmaps between the samples of each morph showed that 10k and 25k spheroids cluster better together than with 50k or 100k, and vice-versa (Figure 2e, f). These observations were consistent across CFL (Figure 2e) and SFL (Figure 2f) spheroid samples. Therefore, for both SFL and CFL, not only did 3D culturing cells into spheroids change transcriptomic profiles from their 2D state, but it also revealed that these profiles are dependent on seeding densities. Consequently, we grouped 10k and 25k as low-density and 50k and 100k as high-density spheroids for all subsequent differential expression (DE) analyses.

**Figure 2:**
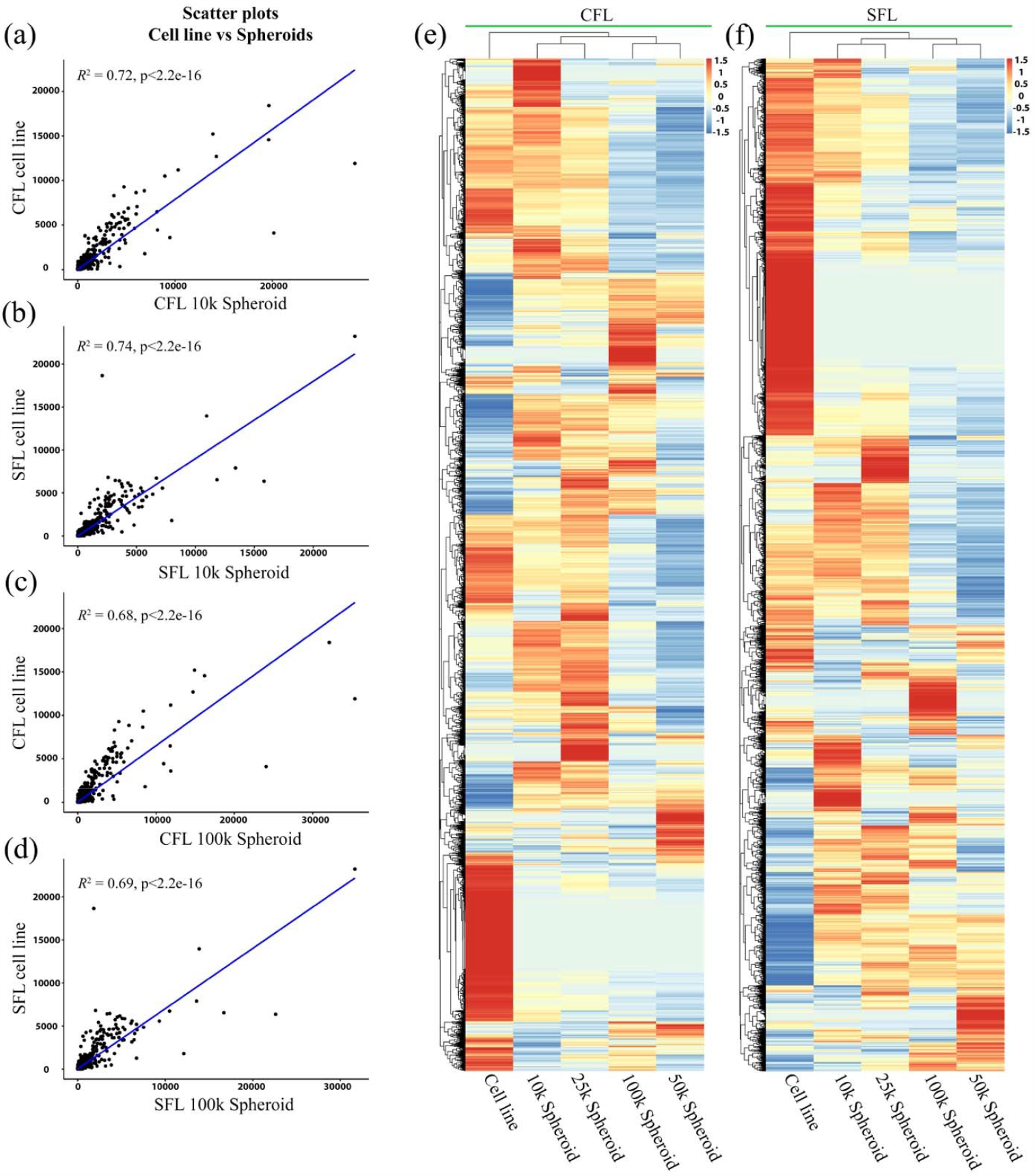
RNA-seq profile of spheroids derived from SFL and CFL cells. (a-d) Scatter plots comparing whole transcriptome of 2D cultures and 3D spheroids. (a) Correlation coefficients - *R*^2^ of CFL spheroids of low-density spheroid of 10k is 0.72. (b) The *R*^2^ of SFL spheroids of low-density spheroid of 10k to 2D cells is 0.74. (c) Correlation coefficients - *R*^2^ of CFL spheroids of high-density spheroid of 100k is 0.68. (d) Correlation coefficients - *R*^2^ of SFL spheroids of high-density spheroid of 100k is 0.69. (e-f) Heatmap clusters of the whole transcriptome between the 2D culture and the different spheroid densities in CFL (e) and SFL (f).

### 3.3. Differential gene expression in low- and high-density spheroids

In order to capture the genes and pathways enriched or repressed upon culturing the cells as spheroids, we performed differential gene expression analysis between low- and high-density spheroids and the cell line. First, we investigated the abundance of the differentially expressed genes across all conditions (Figure 3a).

**Figure 3:**
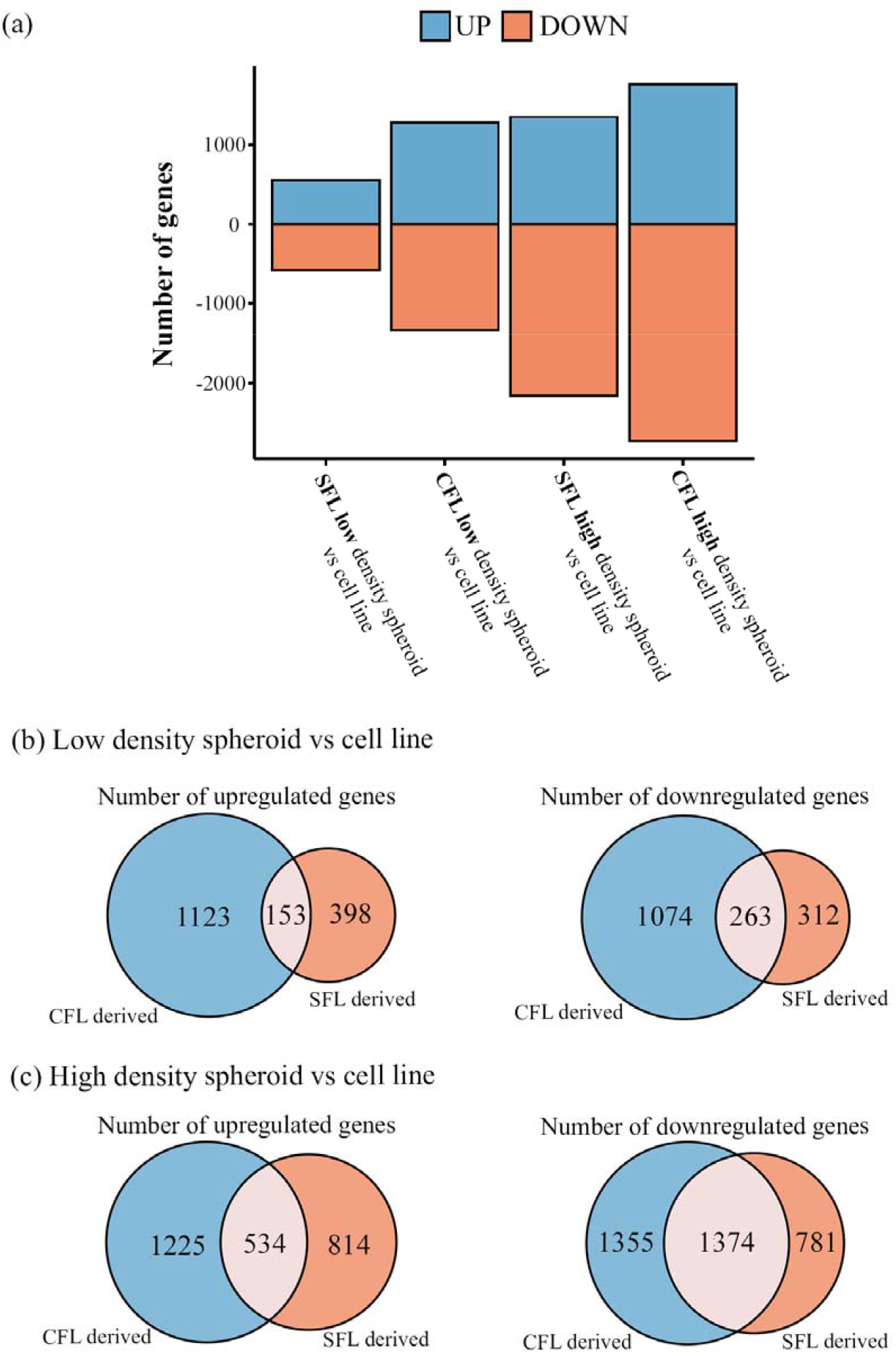
Differential gene expression analysis. (a) Abundance of differentially expressed genes across low and high-density spheroids against 2D culture. (b-c) Overlap of differentially expressed genes between SFL and CFL derived spheroids of low (b) and high density (c) seeding.

When compared to the 2D culture, low-density SFL spheroids showed an upregulation of 551 genes and concomitant downregulation of 575 genes (Figure 3a). CFL spheroids had 1276 genes upregulated, and 1337 which were downregulated (Figure 3a). In the case of high-density seeding, SFL spheroids had 1348 genes upregulated and 2155 genes downregulated with respect to the 2D culture (Figure 3a). The CFL high-density spheroids showed 1759 genes as upregulated and 2729 downregulated (Figure 3a). Taken together this analysis corroborates the findings from the scatter plot analysis by demonstrating that for both SFL and CFL cells, the high-density spheroids exhibit greater changes in expression compared to the ones seeded at lower densities. This effect was greater for CFL than for SFL cells (Figure 3a).

### 3.4. Common response to 3D culture in *Astyanax* liver-derived cells

Next, we identified the pathways associated with the differentially expressed genes. To identify a common response to 3D culturing in the *Astyanax* cells, we looked at common differential expressed genes across low and high-density seedings, spanning both SFL and CFL cells. We identified 153 upregulated genes and 263 downregulated genes across SFL and CFL spheroids of low-density when compared to 2D culture (Figure 3b). In the case of high-density spheroids, we observed 534 commonly upregulated genes and 1374 commonly downregulated genes across SFL and CFL spheroids when compared to the 2D transcriptome (Figure 3c).

Next, we performed Gene Ontology (GO) term analysis. We examined the enriched pathways for both low and high-density spheroids individually and then identified the common ones among the two (Supporting information Figure S3, S4). Among the set of commonly downregulated genes, we identified multiple biological processes enriched. Notably, most of these prominent pathways were associated either directly or indirectly with cell cycle regulation (Supporting information Figure S3). Genes promoting proliferation, such as *etc2, chek1, kif20a* and *plk1* (Lee, Jang, & Lee, 2014; Pabla, Bhatt, & Dong, 2012; Ren et al., 2020; Ulke et al., 2019) were downregulated in both SFL and CFL 3D spheroid samples compared to 2D cultures (Figure 4a, b). Importantly, cell cycle repression is one of the well-known hallmarks of establishing 3D cultures across different cell types (Edmondson, Broglie, Adcock, & Yang, 2014; Lagies et al., 2020; Rybkowska et al., 2023), suggesting that both SFL and CFL cells follow general 3D culturing trends observed in other established cells of different origins.

**Figure 4:**
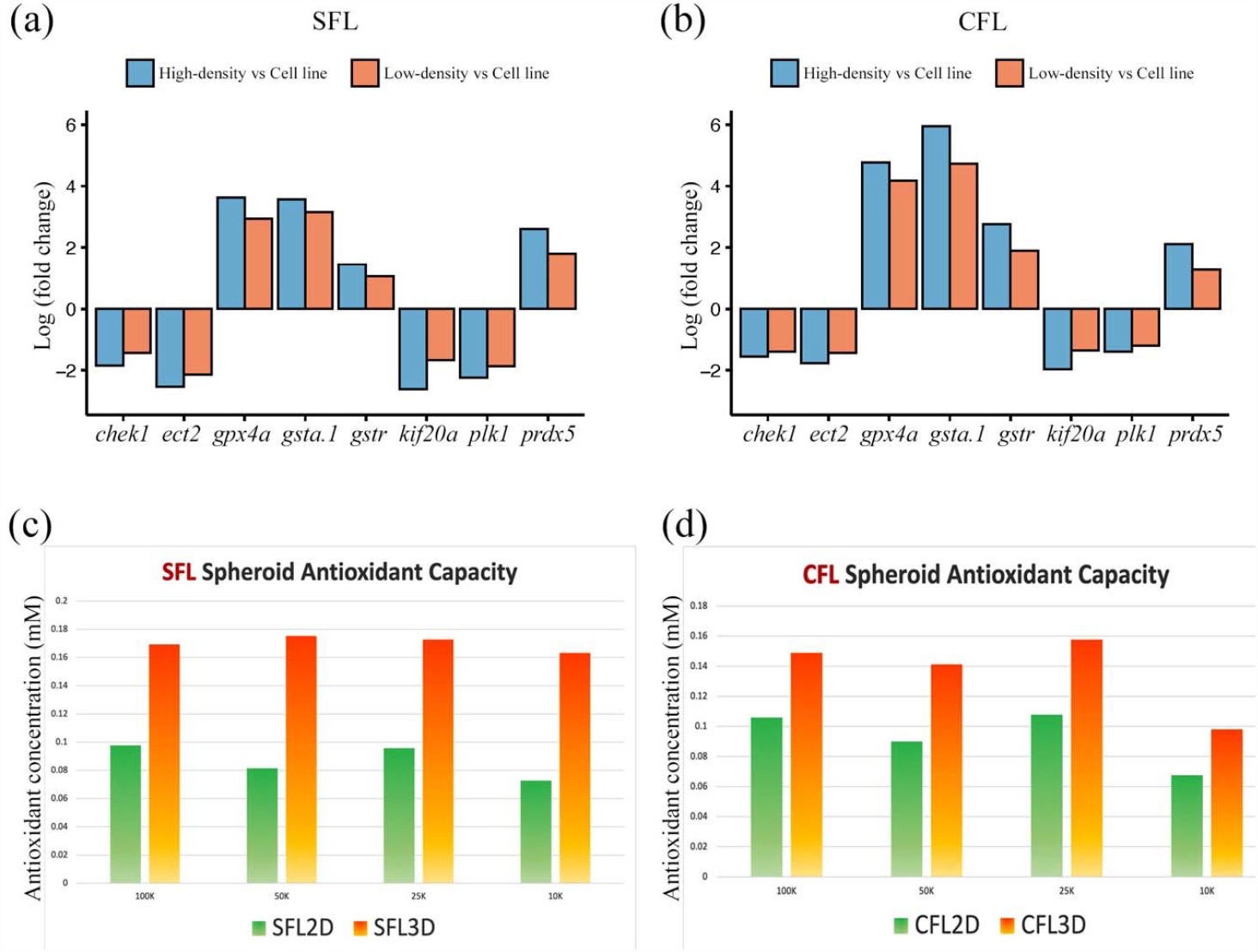
Common response to being 3D cultured. (a-b) Common differential expressed genes across SFL (a) and CFL (b) 3D. Y-axis shows log(fold-change) calculated after averaging TPM values across different seeding densities. (c-d) Comparison between antioxidative capacity of 3D spheroids and 2D cultures of SFL and CFL cells. Both SFL (c) and CFL (d) cells cultured in different cell seeding densities show elevated levels of antioxidative capacity when grown into spheroids (orange bars) with respect to 2D cultures of same seeding density (green bars). The actual values are listed in Supplementary Table S1.

To explore pathways that were induced through 3D culture, we investigated the common upregulated pathways of SFL and CFL spheroids. Using GO term analysis, we identified a list of molecular functions enriched in low and high-density *Astyanax* spheroids (Supporting information Figure S4). We noticed an enrichment for transporter activities such as ‘*inorganic cation transmembrane transporter activity*’, ‘*metal ion transmembrane transporter activity*’ and ‘*cation transmembrane transporter activity*’. Also, we observed an enrichment of peptidase-related terms such as ‘*peptidase activity’* and ‘*exopeptidase activity*’. The analysis also highlighted several antioxidative pathways such as ‘*antioxidant activity*’, ‘*glutathione transferase activity*’, ‘*peroxidase activity*’ and ‘*oxidoreductase activity, acting on peroxide as acceptor*’. Genes like *prdx5, gpx4a*, gstr and *gsta*.*1* which are known to be involved in antioxidative pathways (Sharma, Yang, Sharma, Awasthi, & Awasthi, 2004; Walbrecq et al., 2015; Yant et al., 2003) were upregulated in 3D spheroids (Figure 4a, b). As antioxidants regulating oxidative stress as well as peptidase activity represent essential aspects of liver metabolism (Cichoz-Lach & Michalak, 2014; Dominguez-Vias, Segarra, Ramirez-Sanchez, & Prieto, 2020; Medley et al., 2022; Mooli, Mukhi, & Ramakrishnan, 2022), this data suggests that culturing the liver-derived cells in 3D increases their ability to perform liver metabolic functions. To test this, we performed an antioxidant assay to measure the total antioxidant capacity of the spheroids and compared it with that of the 2D cultured cells (Standard Curve: Supporting information Figure S5). In line with the transcriptomic data, we observed that the spheroids derived from both SFL (Figure 4c) and CFL (Figure 4d) exhibited an enhanced antioxidative capacity compared to their 2D counterpart with similar cell seeding numbers (Supporting information Table S1).

### 3.5. Morph-specific differentially expressed genes in spheroids

Lastly, we aimed to understand the morph-specific transcriptomic responses to 3D culturing. Out of the 1123 and 1225 genes exclusively upregulated (Figure 3b, c) in CFL spheroids, 761 were common in all seeding densities (Figure 5a). Similarly, there were 294 genes exclusively upregulated only in SFL spheroids irrespective of the culture densities (Figure 5b). A similar trend was observed in the commonly downregulated genes, with 592 and 172 in CFL and SFL spheroids, respectively (Figure 5a, b). GO term analysis of the 761 upregulated genes exclusive to CFL spheroids (Figure 5a) revealed biological processes enriched related to xenobiotics, like ‘*response to xenobiotic stimulus*’ and ‘cellular *response to xenobiotic stimulus*’ (Figure 5c) driven by genes such as *sultb16, cyb5a*, and *ache*. Interestingly, the cave morph of *Astyanax* is already known to be more resilient to xenobiotic exposure from the anesthetic MS222 (Bilandzija, Abraham, Ma, Renner, & Jeffery, 2018). In contrast, SFL-specific upregulated genes were enriched for immune responses with GO-terms such as ‘*activation of immune response*’ and ‘*humoral immune response*’ (Figure 5d). This observation also echoes the biology of these morphs, as the surface fish are known to possess enhanced immune responses compared to their cave counterpart (Peuss et al., 2020).

**Figure 5:**
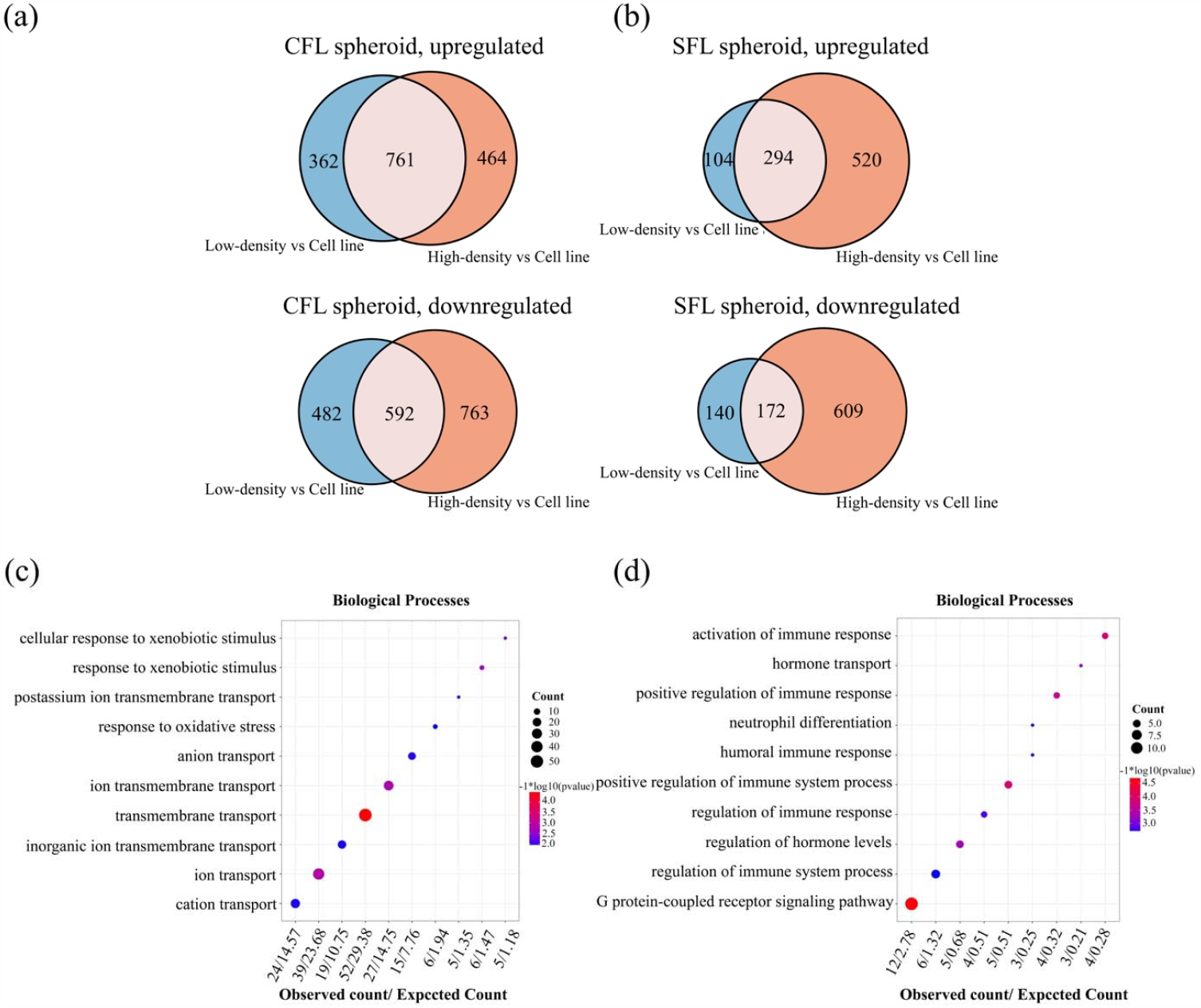
Morph-specific response across SFL and CFL spheroids. (a-b) Morph-specific genes, with overlaps highlighting the common CFL-only (a) and SFL-only (b) differentially expressed genes shared among the different seeding densities. Top panel in both depicts upregulation and bottom panel depicts downregulation. (c) Biological processes enriched for genes exclusively upregulated in CFL spheroids when compared to 2D culture, across varying seeding densities. (d) Biological processes enriched for genes exclusively upregulated in SFL spheroids compared to 2D culture, irrespective of seeding densities.

## 4. Discussion

In this study, we successfully developed and characterized 3D spheroids of *Astyanax* surface and cavefish liver-derived cells. As a common response to 3D culturing, we observed inhibition of cell proliferation markers, which strongly suggests that the liver-derived cells growing in 3D are sensitive to contact inhibition. Importantly we found that many liver-associated metabolic pathways were enriched in both SFL and CFL 3D spheroids, not represented under 2D conditions. Exploring these pathways, we found that spheroids showed enhanced antioxidative capacity *in vitro*. However, we did not detect significant differences between SFL and CFL spheroids in their levels of antioxidative capacity. This is in contrast to Medley et al., which showed an increased antioxidant response in cavefish livers compared to surface fish livers (Medley et al., 2022). However, it is important to note that these experiments were performed *in vivo* in which the cavefish livers are exposed to large amounts of body and liver fat accumulation. As such, it is plausible that under unstressed conditions antioxidant capacities are similar, while under various stresses, the responsive capacities might differ between the morphs. Future experiments will be needed to disentangle this possibility.

Although these 3D enriched pathways provide a more representative metabolic model *in vitro*, reproducing an actual liver in a dish with all its metabolic complexities, is a difficult task. To achieve this goal, we are currently working to develop *Astyanax* liver-derived organoids, by co-culturing different liver-derived cells in 3D. Our present study, essentially being the first example of successful 3D culture of *Astyanax* liver-derived cells, is a vital step in augmenting future research towards *in vitro* studies of *Astyanax mexicanus* liver as well as metabolism.

## Acknowledgements

The authors would like to thank Fengyan Deng, previously from Stowers Histology and presently in Cells, Tissues, and Organoids Center, for her help in sectioning, staining, and imaging experiments. Authors would also like to thank Amanda Lawlor from Stowers Sequencing and Discovery Genomics team for help regarding Library Preparation and Sequencing. This work is supported by institutional funding, and NIH New Innovator Award 1DP2AG071466-01, NIH R24OD030214 and NSF EDGE Award 1923372.

## Conflict of Interest

The authors declare no conflict of interests.

## Data Availability Statement

Original data underlying this manuscript can be accessed from the Stowers Original Data Repository at http://www.stowers.org/research/publications/libpb-2401. The RNA-seq datasets have been uploaded to GEO database under the accession number GSE234212.

## Author contribution

T.B. - Conceptualization, Investigation, Analysis, Visualization, Original draft preparation; N. Rajendran – Conceptualization, Investigation, Visualization; H.H. - Analysis, Visualization, Data Curation; C.Z. - Conceptualization; N. Rohner – Conceptualization, Original draft preparation, Funding acquisition.

## Notes

### Competing Interest Statement

The authors have declared no competing interest.

