## Supporting_information for "3D spheroid culturing of *Astyanax mexicanus* liver-derived cell lines recapitulates distinct transcriptomic and metabolic states of *in vivo* tissue environment"

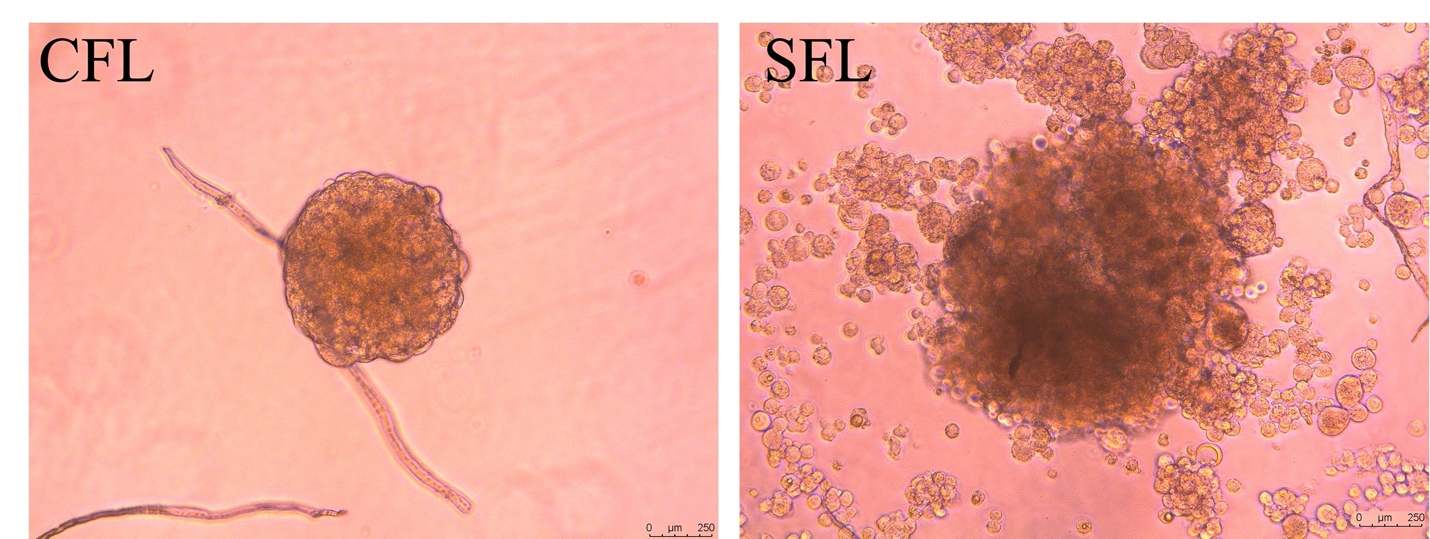

**Figure S1**: Spheroids from CFL and SFL liver-derived cells of *Astyanax* *mexicanus* after 28days in culture.

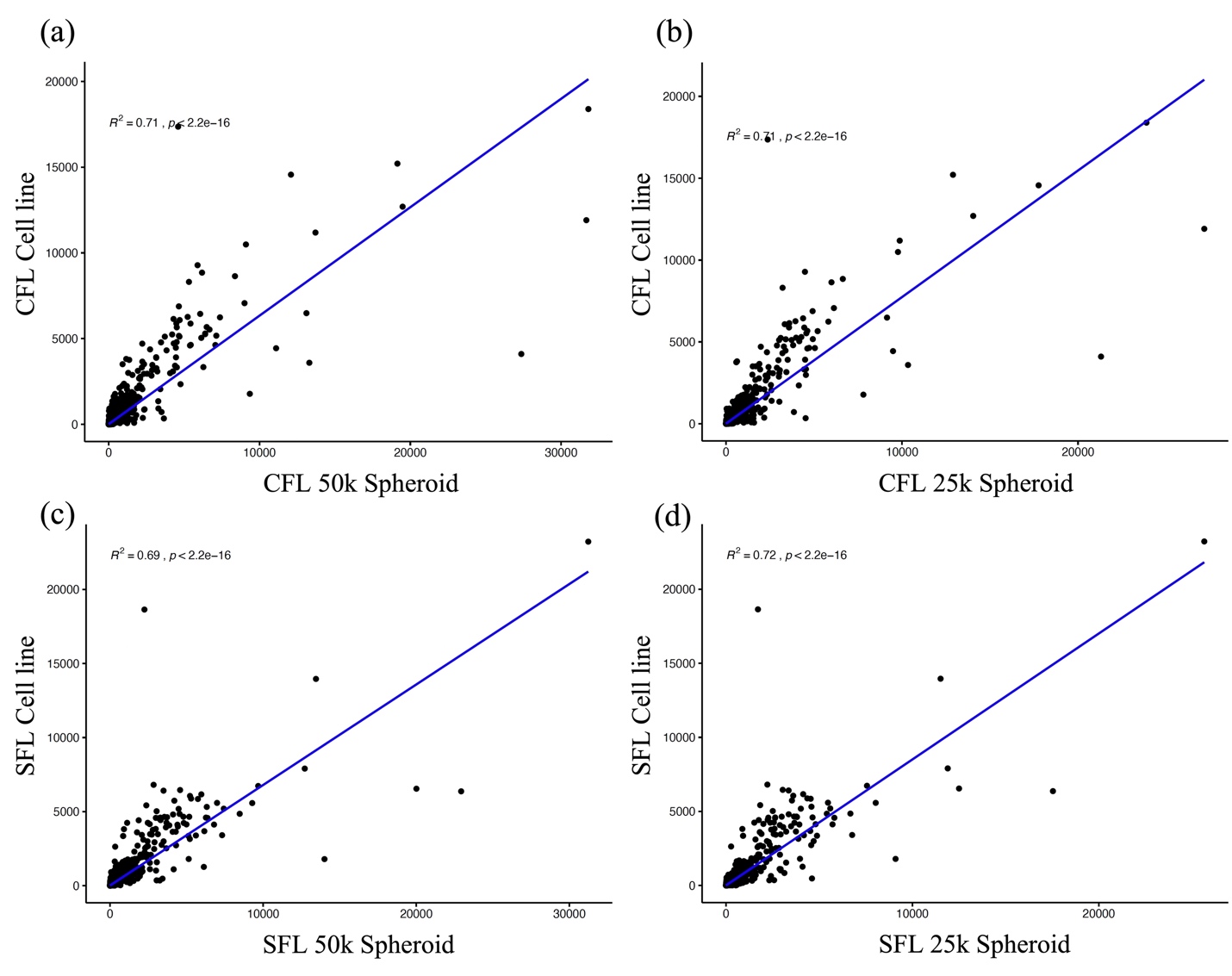

**Figure S2**: Rest of the scatter plots (see Figure 2a -d) comparing transcriptome of 2D monolayer culture and 3D spheroids. (a) Correlation coefficients - *R*^2^ of CFL spheroids of high-density spheroid of 50k is 0.71. (b) The *R*^2^ of CFL spheroids of low-density spheroid of 25k to 2D cells is also 0.71. (c) Correlation coefficients - *R*^2^ of SFL spheroids of high-density spheroid of 50k is 0.69. (d) Correlation coefficients - *R*^2^ of SFL spheroids of low-density spheroid of 25k is 0.72.

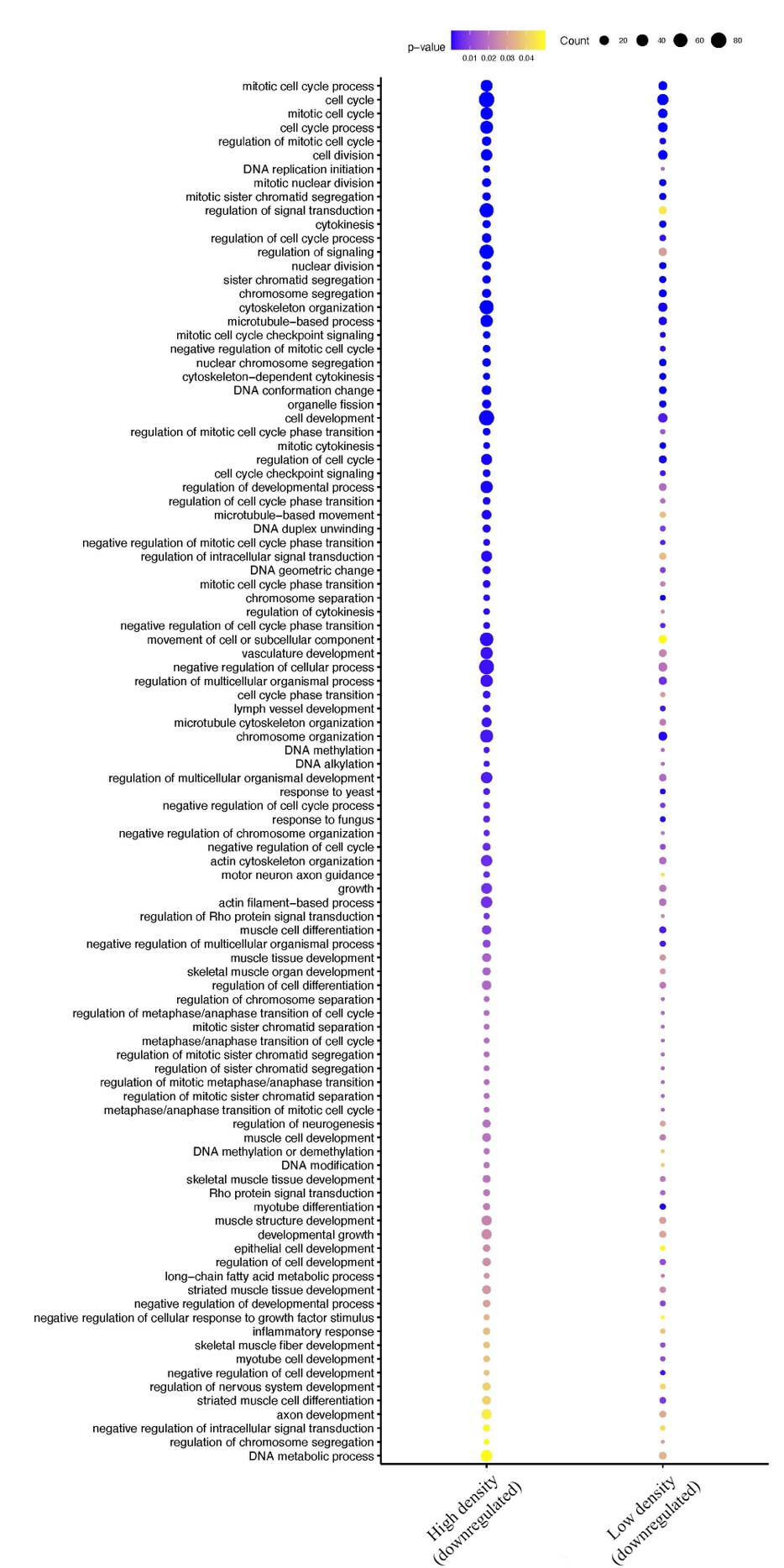

**Figure S3**: Common pathways enriched for downregulated genes in both low and high-density spheroids across SFL and CFL 3D cultures. Majority of the pathways associated with cell cycle and cell division.

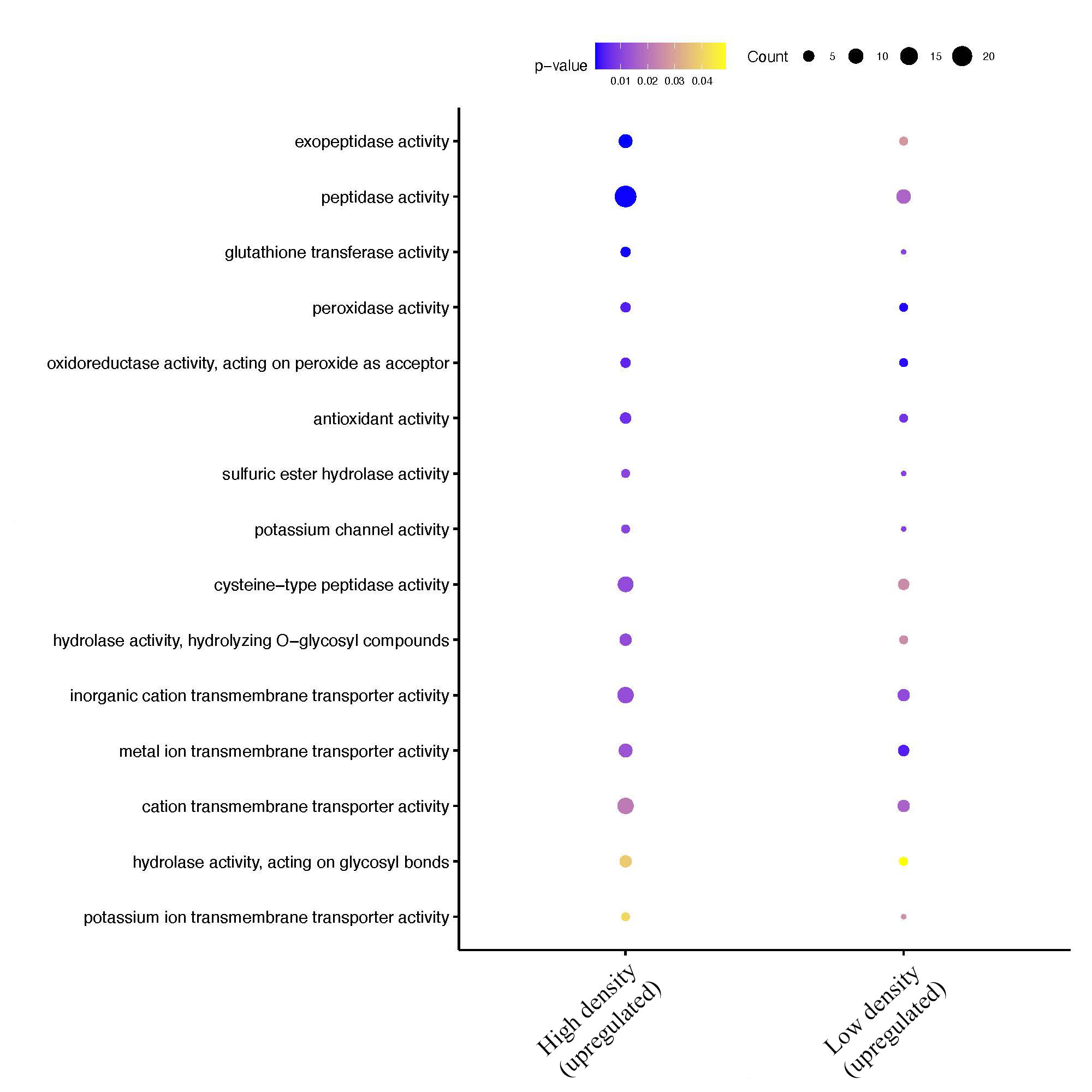

**Figure S4**: Common pathways enriched for upregulated genes in both low and high-density spheroids across SFL and CFL 3D cultures. Majority of the pathways associated with cell cycle and cell division. Antioxidant associated pathways among the top 6 enriched pathways were ‘*antioxidant activity*’, ‘*glutathione transferase activity*’, ‘*peroxidase activity*’ and ‘*oxidoreductase activity, acting on peroxide as acceptor*’.

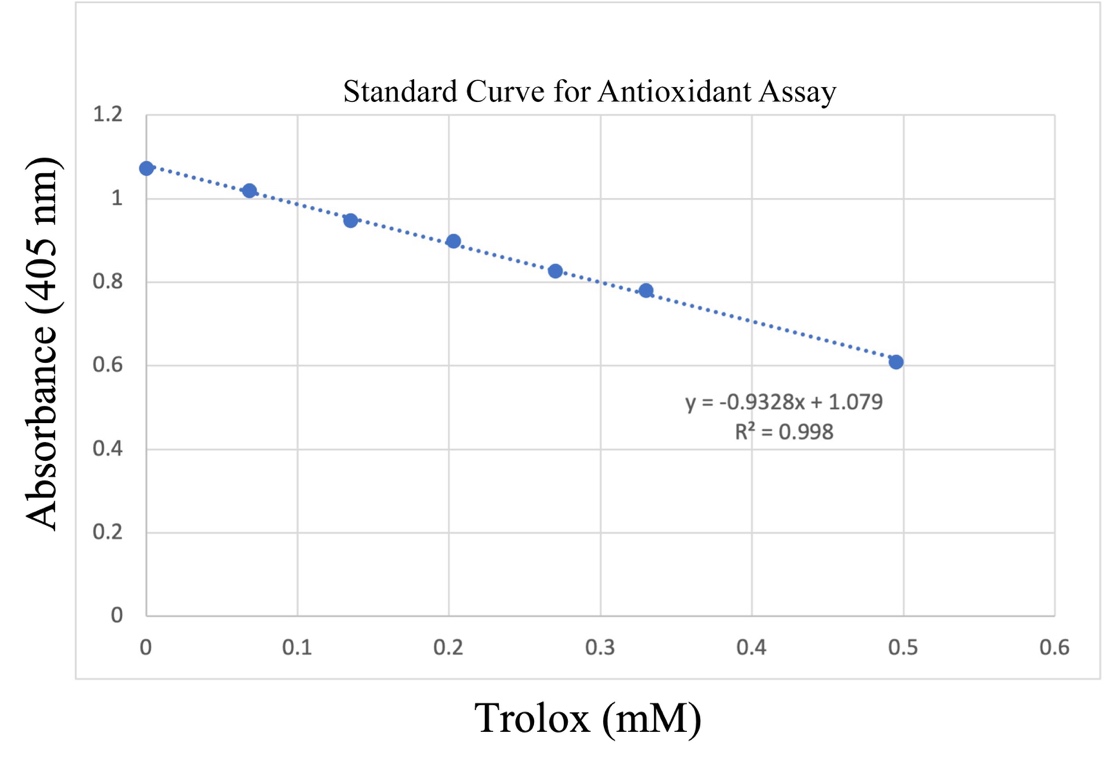

**Figure S5**: The plot depicts the standard curve for measuring antioxidant concentration using colorimetric Antioxidant Assay Kit from Cayman Chemical (Item No. 709001).

|  | SFL2D | SFL3D | CFL2D | CFL3D |
| --- | --- | --- | --- | --- |
| 100K | 0.09766295 | 0.1692753 | 0.10591767 | 0.14890652 |
| 50K | 0.08152873 | 0.17527873 | 0.08999786 | 0.14118782 |
| 25K | 0.09578688 | 0.17270583 | 0.10774014 | 0.15769726 |
| 10K | 0.07268439 | 0.16321827 | 0.06753859 | 0.09803816 |

**Table S1**: Antioxidant concentration (mM) measured across varying cell seeding densities (10k, 25k, 50k and 100k) of regular 2D and 3D spheroid culture of SFL and CFL cells. See Figure 4c, d.
